# The graphene-based affinity cryo-EM grid for the endogenous protein structure determination

**DOI:** 10.1101/2025.02.22.638683

**Authors:** Sojin An, Eungjin Ahn, Tyler Koo, Soyoung Park, Boeon Suh, Krishna P. Rengasamy, Gaocong Lyu, Cheal Kim, Byungchul Kim, Hanseong Kim, Sangho Park, Dongyan Tan, Uhn-Soo Cho

**Author notes:** These authors contributed equally to this work (Sojin An, Eungjin Ahn, Tyler Koo).

## Abstract

Following recent advancements in cryo-electron microscopy (cryo-EM) instrumentation and software algorithms, the next bottleneck in achieving high-resolution cryo-EM structures arises from sample preparation. To overcome this, we developed a graphene-based affinity cryo-EM grid, the Graffendor (GFD) grid, to target low-abundance endogenous protein complexes. To maintain grid quality and consistency within a single batch of 36 grids, we established a one-step crosslinking batch-production method using genetically modified ALFA nanobody as affinity probe (GFD-A grid). Using low concentrations of β-galactosidase-2xALFA, we demonstrated the GFD-A grid’s efficiency in capturing tagged proteins and resolving its cryo-EM structure at 2.71 Å. To test its application for endogenous proteins, we engineered yeast cells with a C-terminal tandem affinity tag (3xALFA-Tev-3xFlag: ATF) at Pop6, a shared component of RNase MRP and RNase P. Cryo-EM structures of RNase MRP and RNase P were resolved at 3.3 Å and 3.0 Å from cell lysates, and 3.6 Å and 3.9 Å from anti-flag elution, respectively. Notably, additional densities were observed in the structures obtained from cell lysates, which were absent in those from the anti-FLAG eluate. These findings establish the GFD-A grid as a robust platform for investigating endogenous proteins, capable of capturing transient interactions and enhancing the resolution of challenging cryo-EM structures with greater efficiency.

## INTRODUCTION

Single-particle cryo-EM has become increasingly significant in structural biology due to its ability to preserve native and dynamic states and to facilitate drug discovery through high-resolution visualization of biomolecular structures^1–3^. Recent technical advancements in cryo-EM equipment, such as the cold field emission gun, aberration corrector, and next-generation direct electron detector, along with software algorithm improvements, have further accelerated this trend^2,4,5^. However, progress in sample and grid preparation steps has been comparatively sluggish and is currently the main bottleneck in high-resolution cryo-EM structure determination^6–12^.

Cryo-EM specimens for single particle analysis can be prepared using either recombinant or endogenous proteins. Recombinant proteins have made significant contributions to structural biology, as many target proteins have been successfully over-expressed and purified using the recombinant expression system^13–15^. However, the recombinant system has limitations in assembling large, multi-subunit complexes due to difficulties in expressing and purifying all components recombinantly and assembling them with proper stoichiometry. As a result, some of cryo-EM structures of large, multi-subunit protein complexes, such as ribosomes and RNA polymerases, have been determined by directly isolating endogenous protein complexes from cells^16–20^. This approach is particularly effective for naturally abundant endogenous protein complexes, as they are relatively easy to isolate and ensure proper folding and assembly in a biologically active conformation. Unfortunately, the same approach is not suitable for most endogenous proteins due to their low copy numbers inside cells. To isolate enough endogenous proteins for cryo-EM studies, a large-scale cell culture and purification system must be established (around 10 ∼ 100 liters for yeasts), making it challenging to explore the structural studies of many important multi-subunit protein complexes in cells^21,22^. This challenge is particularly exacerbated when dealing with limited specimen quantities, which we cannot easily scale up–such as disease variants of endogenous target proteins obtained from human patients or animal models of human diseases^23^. Utilizing these restricted quantities of endogenous proteins for structural studies with current technology is nearly infeasible.

To overcome this challenge, we have developed a cryo-EM grid tool for determining the cryo-EM structures of low-abundant, endogenous protein complexes. The tool is based on a graphene-based affinity cryo-EM grid, which we refer to as the Graffendor Grid (<u>Gr</u>aphene-based <u>aff</u>inity cryo-EM grid for the <u>endo</u>genous protein complex; GFD Grid). Particularly, we paid attention on specimen loss during the blotting step. After protein purification, most of purified proteins (more than 99%) are absorbed into the filter paper during the blotting step. Minimizing specimen loss by attracting proteins on the grid surface before blotting would reduce sample demand dramatically.

Developing the affinity grid has been a hot topic in the field, and several earlier attempts have been reported, including the nickel-nitrilotriacetic acid (Ni-NTA) functionalized lipid monolayer grid^24^, the antibody-decorated affinity grid^25,26^, the Ni-NTA coated graphene-oxide grid^27^, the monolayer crystal of streptavidin-coated affinity grid^28^, and the SpyCatcher/SpyTag-mediated graphene-oxide grid^29^. While these pioneering studies have demonstrated the potential impact of the affinity grid in the cryo-EM field, its practical implementation has been limited probably by the inconsistent quality control of the affinity grid arising from intricate multi-step procedures and one-by-one manufacturing approach. The Graffendor grid we developed here addresses this limitation by incorporating a single-step crosslinking procedure of the affinity probe (modified NbALFA) onto the graphene grid substrate (Fig. 1a) and facilitates the simultaneous production of 36 affinity grids in a single batch^30^, thereby ensuring consistent quality across all grids within the same batch. This ease of use and efficient workflow make the Graffendor grid an attractive tool for studying challenging endogenous protein structures and addressing key questions in protein function and regulation via structures.

**Figure 1.**
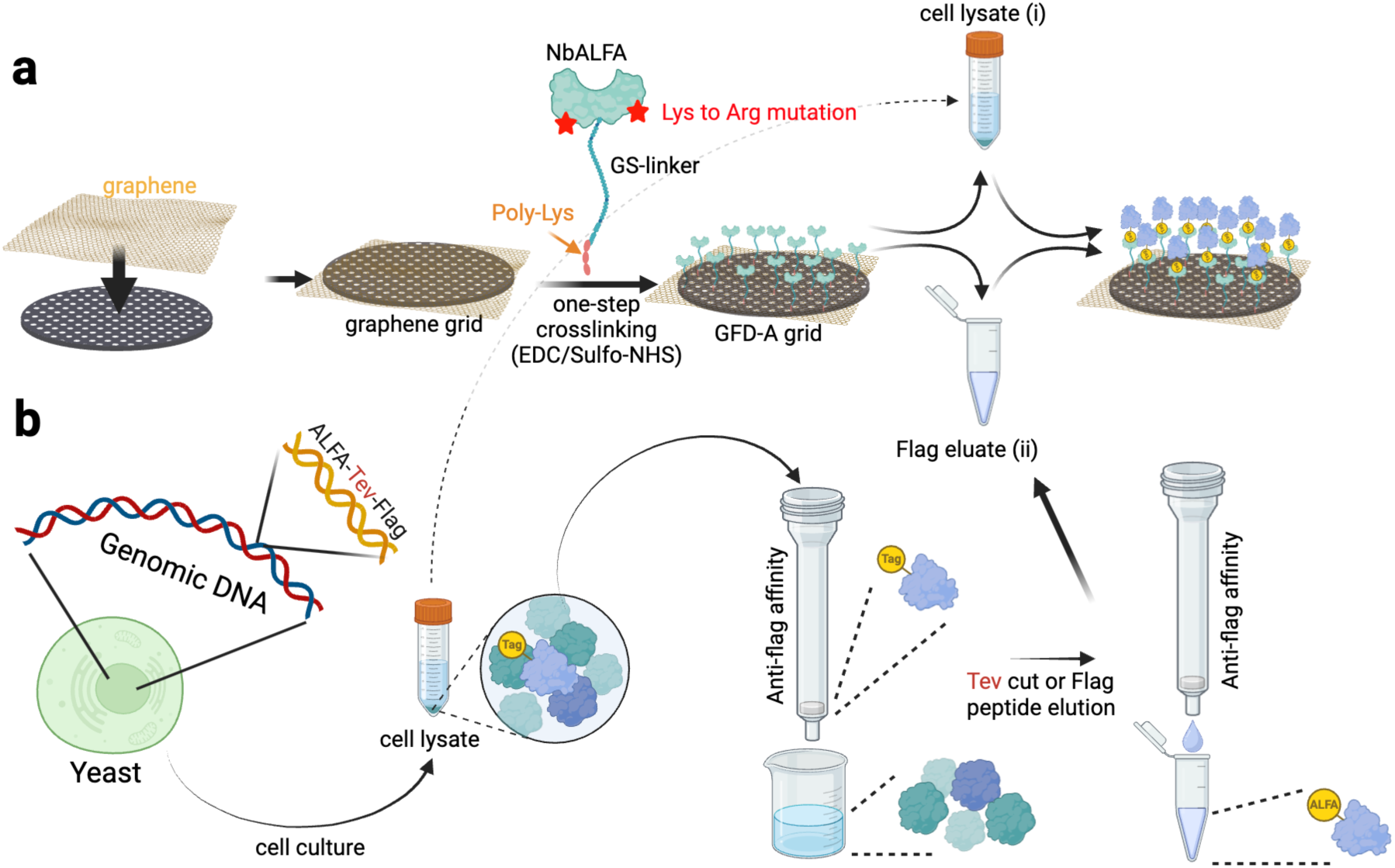
**a, b** Overall scheme of the GFD-A grid production (a) and incubation of the GFD-A grid with either cell lysates (i) or the affinity eluate (ii) of TAP-tagged endogenous proteins from yeast (b) for cryo-EM studies (created by biorender.com).

In this report, we have developed a graphene-based affinity grid using the modified ALFA nanobody (NbALFA) as an affinity probe, which we refer to as the GFD-A type grid (Graffendor-ALFA; Fig. 1a)^30^. We have also generated stable yeast strains containing a tandem affinity purification (TAP)-tag (ATF; 3x <u>A</u>LFA tag–<u>T</u>ev site–3x <u>F</u>lag tag) at the C-terminal end of the target gene using homologous recombination^31^ (Fig. 1b). After cell culture and lysis, the target protein, which is mostly a low abundant endogenous protein, can be purified using anti-flag affinity chromatography. The GFD-A grid is either directly incubated in cell lysates (i) or immersed in the affinity eluate (ii) to capture purified target proteins on the grid. The GFD-A grid is washed, plunge-frozen, and subsequently used for imaging in cryo-EM.

## Results

### Making the ALFA-nanobody attached Graffendor grid (GFD-A grid)

In a prior study, we developed a robust and reproducible method to produce graphene grids in large quantities, yielding 36 grids per batch^30^. These graphene grids underwent oxidation via a brief plasma treatment, introducing functional groups such as hydroxyl, epoxyl, and carboxyl groups to the graphene layer^32,33^. The oxidized surface was then chemically activated using EDC (1-ethyl-3-[3-dimethylaminopropyl]carbodiimide hydrochloride) and Sulfo-NHS (N-hydroxysulfosuccinimide) to enable crosslinking with amine groups^34^. The ALFA-tag-specific nanobody (NbALFA; 13.6 kDa) has a strong affinity (∼26 pM) for the ALFA-tag (15 amino acids; PSRLEEELRRRLTEP), making it ideal for our application^35^. NbALFA was genetically modified for optimal crosslinking on the graphene surface by (1) mutating two lysine residues (K80R and K90R) to arginine to prevent unintended direct crosslinking of NbALFA to the activated graphene grid, (2) extending the C-terminal end with a 32-residue GS-linker to provide flexibility (∼100 Å from the graphene surface) and reduce preferred particle orientation, and (3) adding five C-terminal poly-lysine residues for efficient chemical crosslinking with the EDC/Sulfo-NHS activated graphene surface (Fig. 1a and Suppl. Fig. 1a). The modified NbALFA (∼2 μM) was incubated with the EDC/Sulfo-NHS activated graphene grid, allowing crosslinking via its C-terminal poly-lysine residues. This process resulted in the creation of the NbALFA-attached Graffendor grid (GFD-A grid). The completed GFD-A grid was subsequently transferred to a storage buffer, making it ready for specimen treatment and further experimental applications (Suppl. Fig. 1b).

### Specificity of the GFD-A grid toward purified β-galactosidase-2xALFA

A functional affinity grid requires two essential properties: (1) selectivity, achieved through specific interactions between the target protein and a probe, and (2) the ability to attract and populate particles on the grid. To evaluate the functionality of the GFD-A grid, we cloned and purified β-galactosidase with a 2xALFA tag at its C-terminus (Suppl. Fig. 2a). An amylose pull-down assay using maltose-binding protein (MBP)-tagged modified NbALFA confirmed the successful capture of β-galactosidase-2xALFA by modified NbALFA (Suppl. Fig. 2b). To test the crosslinking efficiency of modified NbALFA on the EDC/Sulfo-NHS-activated graphene surface, we introduced an N-terminal cysteine (E2C mutation) and labeled NbALFA with sulfo-Cyanine5 maleimide (Cy5). However, the Cy5-labeled NbALFA formed aggregates, likely due to interference with its internal disulfide bond. To address this issue, we alternatively used an N-terminal cysteine-incorporated modified calmodulin (5 surface Lys residues to Arg; 34 GS linker and 5 poly-lysine residues at the C-terminus) for Cy5 labeling (Suppl. Fig. 2c). Initially, two types of affinity grids were developed: NbALFA-based (GFD-A) and calmodulin-based (GFD-C), but only the GFD-A grid functioned as intended. N-terminal Cy5-labeled modified calmodulin displayed strong fluorescence signals at a wavelength of 670 nm under the Nikon CSU-X1 confocal microscope (Nikon, Japan), confirming successful EDC/Sulfo-NHS-mediated crosslinking (Suppl. Fig. 2d, e). In comparison, Cy5-labeled modified calmodulin without EDC/Sulfo-NHS treatment exhibited minimal fluorescence under identical conditions. This indicates that modified NbALFA, using the same EDC/Sulfo-NHS crosslinking approach, was effectively coated on the graphene surface via chemical crosslinking.

To assess the ability of the GFD-A grid to capture and concentrate low levels of β-galactosidase-2xALFA from solution, we incubated the grid with 500 µl of β-galactosidase-2xALFA at concentrations of 5 µg/ml and 25 µg/ml (Fig. 2a, c). We used purified β-galactosidase-CBP (calmodulin-binding peptide) as a negative control (Fig. 2b, d; Suppl. Fig. 2a). SDS-PAGE analysis showed that β-galactosidase-2xALFA at 5 µg/ml was faintly visible with Coomassie stain but detected using silver stain (Suppl. Fig. 3). Cryo-EM micrographs revealed that β-galactosidase-2xALFA densely populated the GFD-A grid, whereas β-galactosidase-CBP did not. The GFD-A grid incubated with β-galactosidase-CBP appeared similar to a GFD-A grid without any specimen (Suppl. Fig. 4). These results demonstrate the specificity of the GFD-A grid for ALFA-tagged proteins and confirm its effectiveness as an affinity grid, even at low protein concentrations.

**Figure 2.**
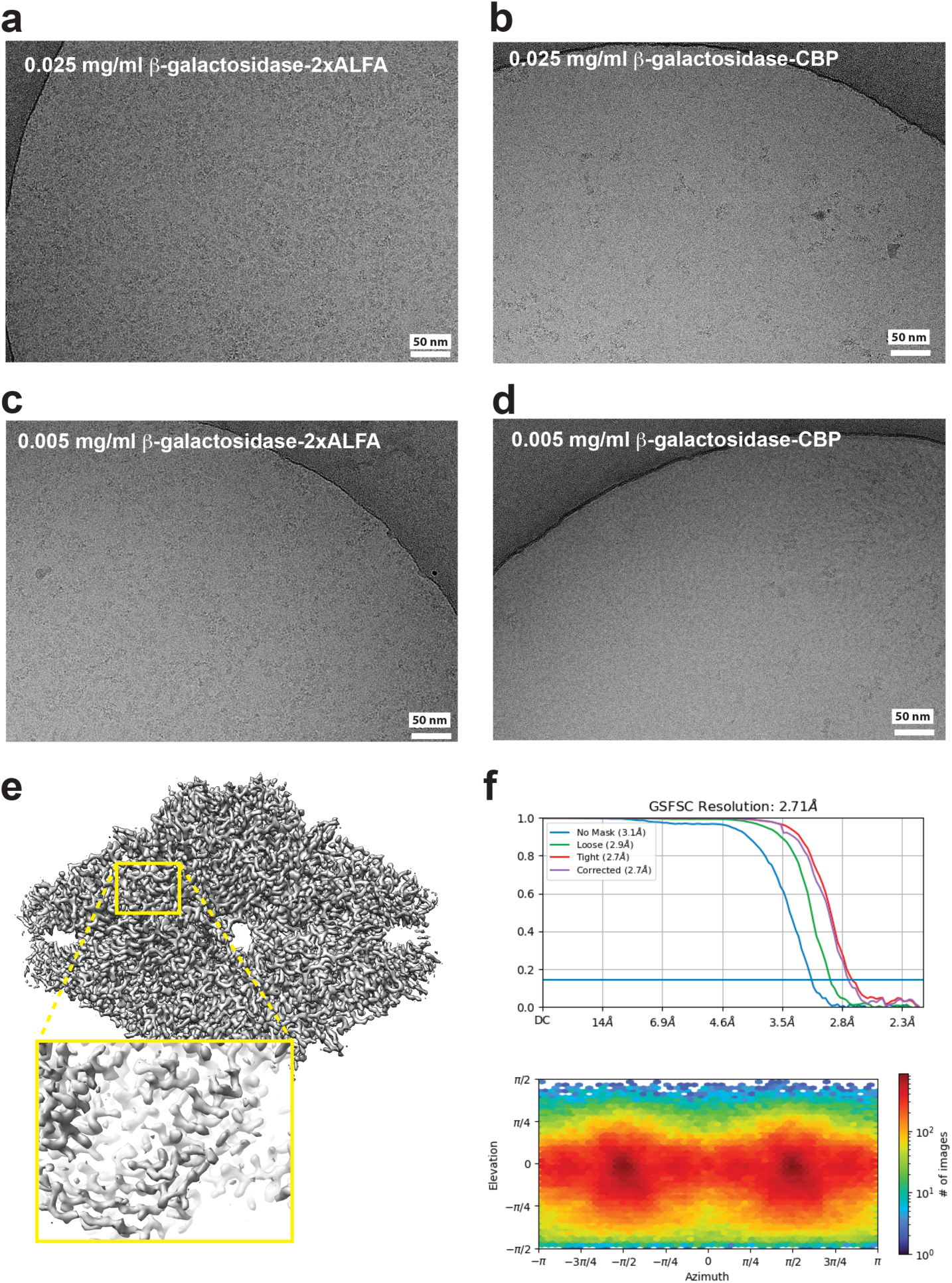
**a,** Micrographic images of the GFD-A grid incubated with 0.025 mg/ml **(a)** and 0.005 mg/ml **(c)** β-galactosidase-2xALFA and 0.025 mg/ml **(b)** and 0.005 mg/ml **(d)** β-galactosidase-CBP (300 KeV Krios, K3 direct electron detector). **e,** Cryo-EM structure of β-galactosidase-2xALFA at 2.7 Å resolution (1,761 micrographs; 1.08 Å/px; 66 e^-^/A^2^/sec; final 336K particles). The yellow square box is the zoom-in view of the final cryo-EM map. **f,** the FSC curve and preferred particle distribution adapted from cryosparc.

To comprehensively evaluate the performance of the GFD-A grid, we used a GFD-A grid bound with 25 µg/ml β-galactosidase-2xALFA and collected 1,760 microscopic images using a 300 KeV Titan Krios microscope equipped with a K3 Summit direct electron detector and a Gatan image filter (±20 eV). A total of 755K particles were picked using TOPAZ^36^, and the cryo-EM structure of β-galactosidase-2xALFA was determined at 2.71 Å resolution with 336K particles using the cryoSPARC program suite^37^ (Fig. 2e, f). These findings highlight the exceptional specificity and efficiency of the GFD-A grid in enabling high-quality cryo-EM structural analysis.

To evaluate the long-term stability of the GFD-A grid, several grids from the same batch were produced and stored in the cold room for up to 8 months in GFD-A storage buffer (30 mM HEPES, pH 7.0, 150 mM NaCl, 0.1% NP-40, and 0.02% NaN; Suppl. Fig. 1b). The results indicated that the GFD-A grid maintained its binding capacity for at least 8 months under these conditions (Suppl. Fig. 5).

### Monitoring particle distribution of β-galactosidase-2xALFA on the GFD-A grid using in vitro electron cryo-tomography (cryo-ET)

One of the major challenges in cryo-EM sample preparation is the protein denaturation mediated by the air-water interface^12,38^. Using the *in vitro* cryo-ET of the graphene-coated grid, we previously demonstrated that particles adhered to the graphene surface, thereby avoiding exposure to the air-water interface^30^. To visualize particle distribution on the graphene-coated GFD-A grid, we performed tilt series imaging of plunge-frozen samples containing 20 μg/ml β-galactosidase-2xALFA bound to the GFD-A grid. For accurate image alignment during data collection, 10 nm gold nanoparticles were introduced as fiducial markers before plunging. The resulting images revealed dense populations of β-galactosidase-2xALFA particles distributed on both sides of the graphene layer with noticeable separation (Fig. 3a,b). Unlike sMMOH particles on the graphene grid, β-galactosidase-2xALFA particles did not directly contact the graphene layer on either side, indicating that a long GS linker (∼ 100 Å) in the modified NbALFA provided a distance barrier from the graphene surface.

**Figure 3.**
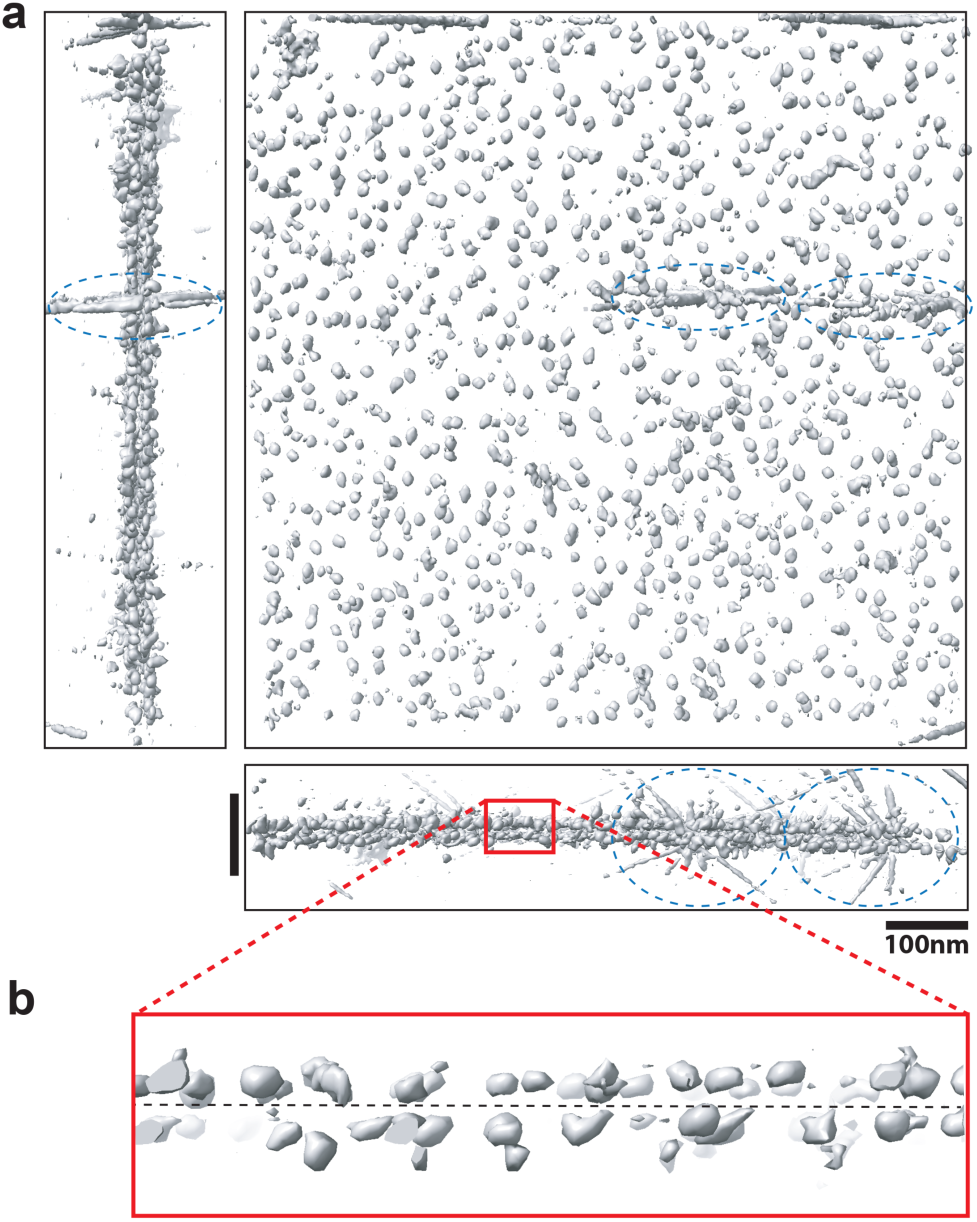
β-galactosidase-ALFA particle distribution on plunge-frozen GFD-A grid. **a,** Top, side, and central views of segmented β-galactosidase-ALFA particles (grey) from a tomograph of the GFD-A grid. The blue dashed circles indicate 10nm gold fiducial marks. **b,** Zoom-in side-view of segmented β-galactosidase-ALFA particles (red box). The black dashed horizontal line indicates the position of the graphene layer.

Furthermore, analysis indicated that the particles maintained an approximate distance of ∼50 nm from the graphene surface (with an ice thickness of ∼100 nm), suggesting that they remained shielded from the air-water interface owing to the presence of NbALFA crosslinked on the graphene layer. Consequently, *in vitro* electron cryo-tomography conducted on the β-galactosidase-2xALFA-bound GFD-A grid illustrated that the particles were positioned in proximity to the graphene layer via modified NbALFA, effectively avoiding exposure to the air-water interface. Additionally, this approach highlighted that the particles maintained a distance from the graphene surface due to the GS linker present in the modified NbALFA.

### Specificity of the GFD-A grid for β-galactosidase-2xALFA expressed in E. coli cell lysate and its suitability for long-term storage

To assess whether the GFD-A grid retains target specificity in the presence of cell lysates, we prepared *E. coli* cell lysates expressing β-galactosidase-2xALFA and incubated them with both a plain graphene grid and the GFD-A grid. As shown in Figure 4, only the GFD-A grid successfully captured β-galactosidase-2xALFA from the lysates, whereas the graphene grid did not. This result confirms the strong specificity of the GFD-A grid for its target proteins, even in complex cellular environments.

**Figure 4.**
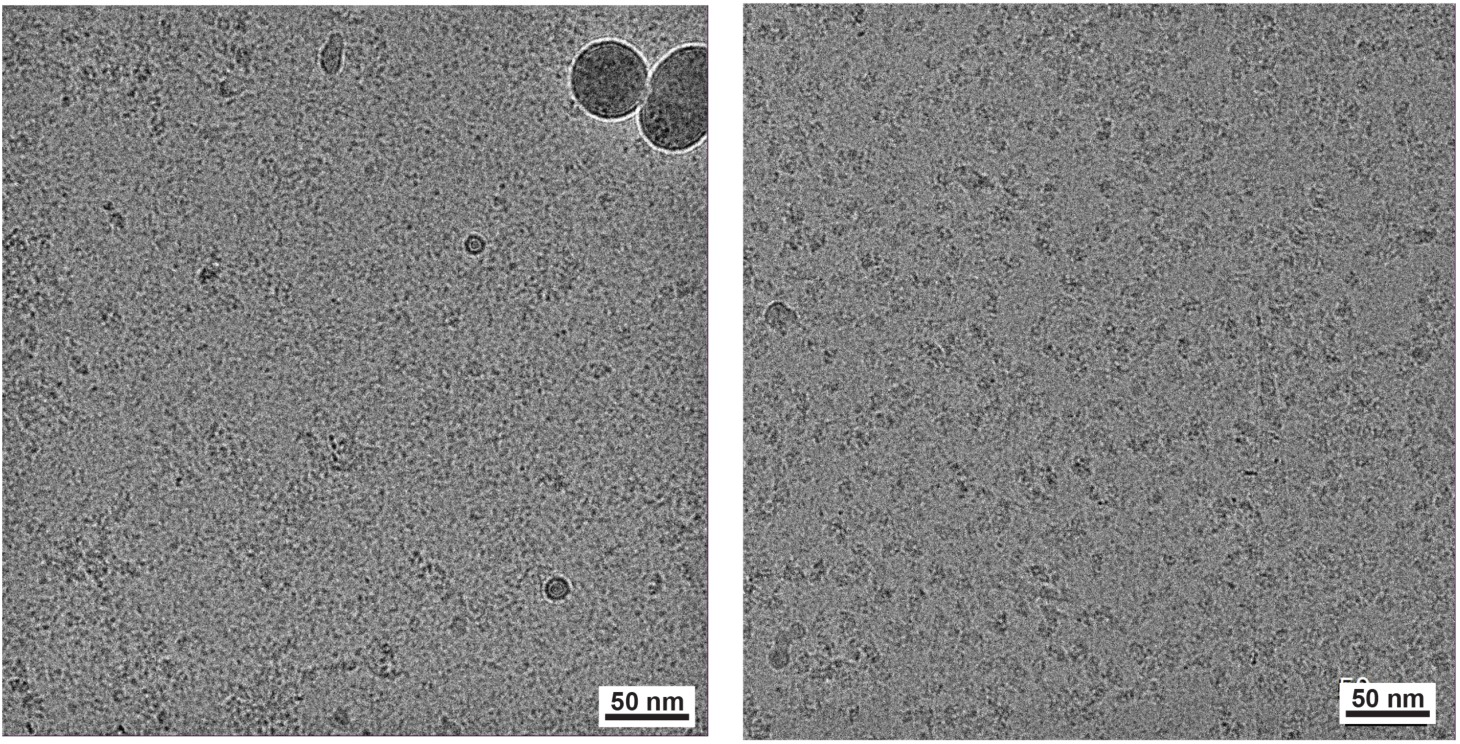
β-galactosidase-2xALFA expressed *E. coli* cell lysates (0.5 ml each) were incubated with the graphene grid (left) and the GFD-A grid (right). Non-specific binding on the graphene grid yet dense β-galactosidase-ALFA particles on the GFD-A grid were observed.

The ideal affinity grid requires to maintain the binding efficiency over time. Therefore, we conducted experiments to evaluate the binding efficiency of the GFD grid after long-term storage for up to eight months. As shown in Supplementary Figure 5, we demonstrated that the GFD grid can be stored in a cold room for at least eight months without any detectable decrease in binding efficiency.

### Cryo-EM structure determination of endogenous RNase P and RNase MRP from budding yeast using the GFD-A grid

To assess the feasibility of the GFD-A grid for capturing and resolving the cryo-EM structures of endogenous protein complexes, we utilized tandem affinity purification (TAP)-tagging (ATF; 3xALFA-TEV-3xFLAG) at the C-terminus of target genes in budding yeast via homologous recombination^31^. As a proof-of-concept, the ATF tag was introduced at the C-terminus of Pop6, a shared component of RNase P^39^ and RNase MRP^40,41^, whose cryo-EM structures have been previously determined at resolutions of 3.5 Å and 3.0 Å, respectively. In these prior studies, large-scale yeast cell cultures (24∼32 liters) were required to isolate sufficient quantities of RNase P and RNase MRP for structural analysis. Using the Pop6-ATF strain, we aimed to evaluate (1) whether atomic-resolution structures of these endogenous complexes can be determined directly from both cell lysate and anti-FLAG eluate after incubation with the GFD-A grid, and (2) whether compositionally heterogeneous complexes can be resolved from a single preparation.

RNase P and RNase MRP in *S. cerevisiae* are ribonucleprotein complexes that share most of protein components (Pop1, Pop3, Pop4, Pop5, Pop6, Pop7, Pop8, and Rpp1), yet contain their unique subunits; Rpr2 in RNase P, and Rmp1/Snm1 in RNase MRP^42^. RNase P is essential for tRNA 5’-end processing^39^, and RNase MRP involves in the maturation of rRNA and metabolism of mRNAs participated in the cell cycle regulation^40,43–45^. Both RNase P and RNase MRP possess catalytic RNA components associated with protein subunits, totaling ∼ 450 kDa.

To evaluate the utility of the GFD-A grid, the TAP tag (ATF) was introduced at the C-terminus of Pop6 (Pop6-ATF) via yeast homologous recombination. One liter of yeast cells expressing endogenous Pop6-ATF was cultured and harvested at an optical density of approximately 2.0. Following cell disruption using a freezer mill, the soluble fractions of the lysates were processed in two ways: (1) directly incubated with the GFD-A grid (Fig. 5a–e; cell lysate/GFD-A) or (2) subjected to anti-FLAG elution after TEV cleavage, followed by incubation with the GFD-A grid (Fig. 5f–j; anti-FLAG eluate/GFD-A). In the cell lysate/GFD-A setup, endogenous RNase P and RNase MRP complexes were selectively captured through Pop6-ATF/NbALFA interactions, effectively isolating these complexes from cellular contaminants (Fig. 5a). For the anti-FLAG eluate/GFD-A approach, RNase P and RNase MRP were enriched through anti-FLAG affinity purification before immobilization on the GFD-A grid (Fig. 5f). Plunge-frozen GFD-A grids from both setups were imaged using a 300 keV Titan Krios microscope equipped with a K3 summit direct electron detector and a Gatan imaging filter (±20 eV). Raw micrographs revealed densely populated particles on the GFD-A grids in both conditions (Figs. 5b, g), confirming the grid’s specificity for ALFA-tagged targets. Using TOPAZ^36^, total 1.17 and 1.10 million particles of RNase P/MRP were picked from 4,924 (cell lysate/GFD-A) and 5,148 (anti-flag eluate/GFD-A) micrographs, respectively (Suppl. Table 1) and underwent 2D classification (Figs. 5c, h). Cryo-EM structural analysis resolved RNase P and RNase MRP from the cell lysate/GFD-A dataset to 3.0 Å (Figs. 5d) and 3.3 Å (Fig. 5e), respectively, and from the anti-FLAG eluate/GFD-A dataset to 3.9 Å (Fig. 5i) and 3.6 Å (Fig. 5j).

**Figure 5.**
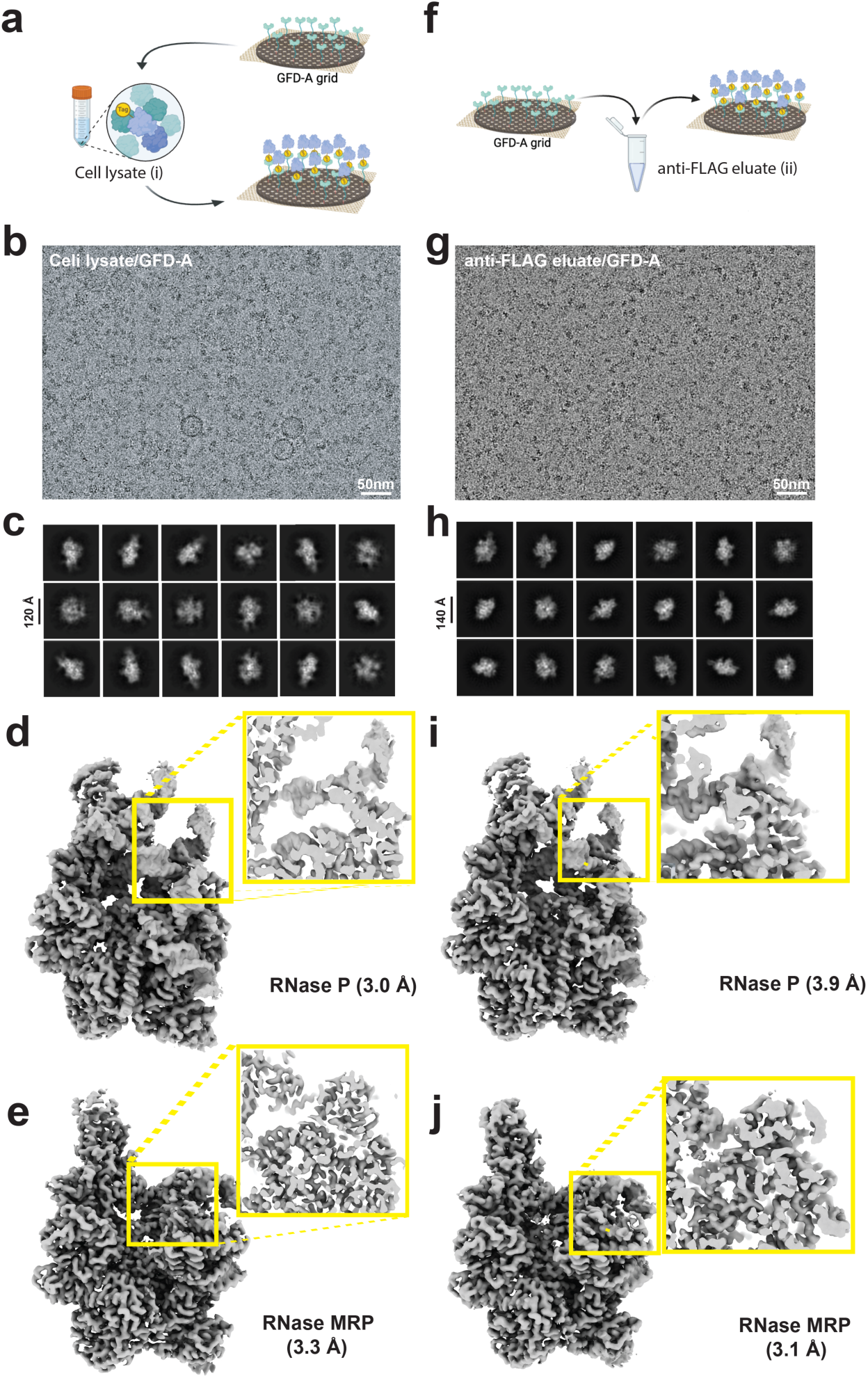
**a-e,** direct incubation of the GFD-A grid in cell lysates (Pop6-ATF) **(a**), representative micrograph (**b**), 2D classification (**c**), and cryo-EM structures of RNase P/MRP (**d-e**) of the cell-lysate/GFD-A grid. **f-j,** incubation of the GFD-A grid in anti-flag eluate **(f)**, representative micrograph (**g**), 2D classification (**h**), and cryo-EM structures of RNase P/MRP (**i-j**) of the anti-flag-eluate/GFD-A grid.

This study demonstrates that the GFD-A grid specifically captures target proteins from both cell lysates and affinity-purified fractions. Furthermore, it highlights the ability of the GFD-A grid to resolve compositionally heterogeneous complexes and determine their cryo-EM structures from a single batch of cell growth.

### Additional cryo-EM densities observed in RNase P/MRP structures from cell lysates

A notable observation when comparing the cryo-EM structures of RNase P/MRP derived from either cell lysate or anti-FLAG eluate was the presence of additional densities in the cell lysate-derived structures at a low contour level (0.002) (Suppl. Fig. 6). These densities were absent in the anti-FLAG eluate-derived RNase P/MRP at the same contour level, suggesting that these extra densities are specifically associated with RNase P/MRP isolated directly from cell lysate. To enhance the quality of these additional cryo-EM densities, we performed multibody refinement. However, due to their low occupancy and likely dynamic nature, we were unable to resolve these densities at near-atomic resolution. Nevertheless, we could determine which RNase P/MRP subunits are linked to these additional features. In RNase P, two distinct extra densities were identified (Fig. 6a): one connected to the N-terminal regions of Rpr2 and Pop3 (Rpr2 residues 1-16, Pop3 residues 1-13; Fig. 6b) and another associated with Rpp1 (Fig. 6c). Interestingly, when these densities were overlaid with the pre-tRNA-bound RNase P structure (PDB ID: 6AH3)^39^, the pre-tRNA fit well between the additional densities and the main body of RNase P. This suggests that these densities may either obstruct pre-tRNA access or contribute to substrate selectivity. For RNase MRP (cell lysate), an additional density was observed adjacent to Snm1/Pop3, extending upward, in contrast to the downward orientation of the corresponding RNase P density linked to Rpr2/Pop3 (Fig. 6e,f). In both cases, the unique subunits of RNase P/MRP (Rpr2 for RNase P and Snm1 for RNase MRP) serve as anchoring points for these extra densities. This observation suggests that the additional densities adjacent to Rpr2 (RNase P) and Snm1 (RNase MRP) may contribute to functional differences between the two ribonucleoprotein complexes. Unfortunately, due to their low occupancy and likely dynamic behavior, we were unable to determine the precise identity of these additional densities.

**Figure 6.**
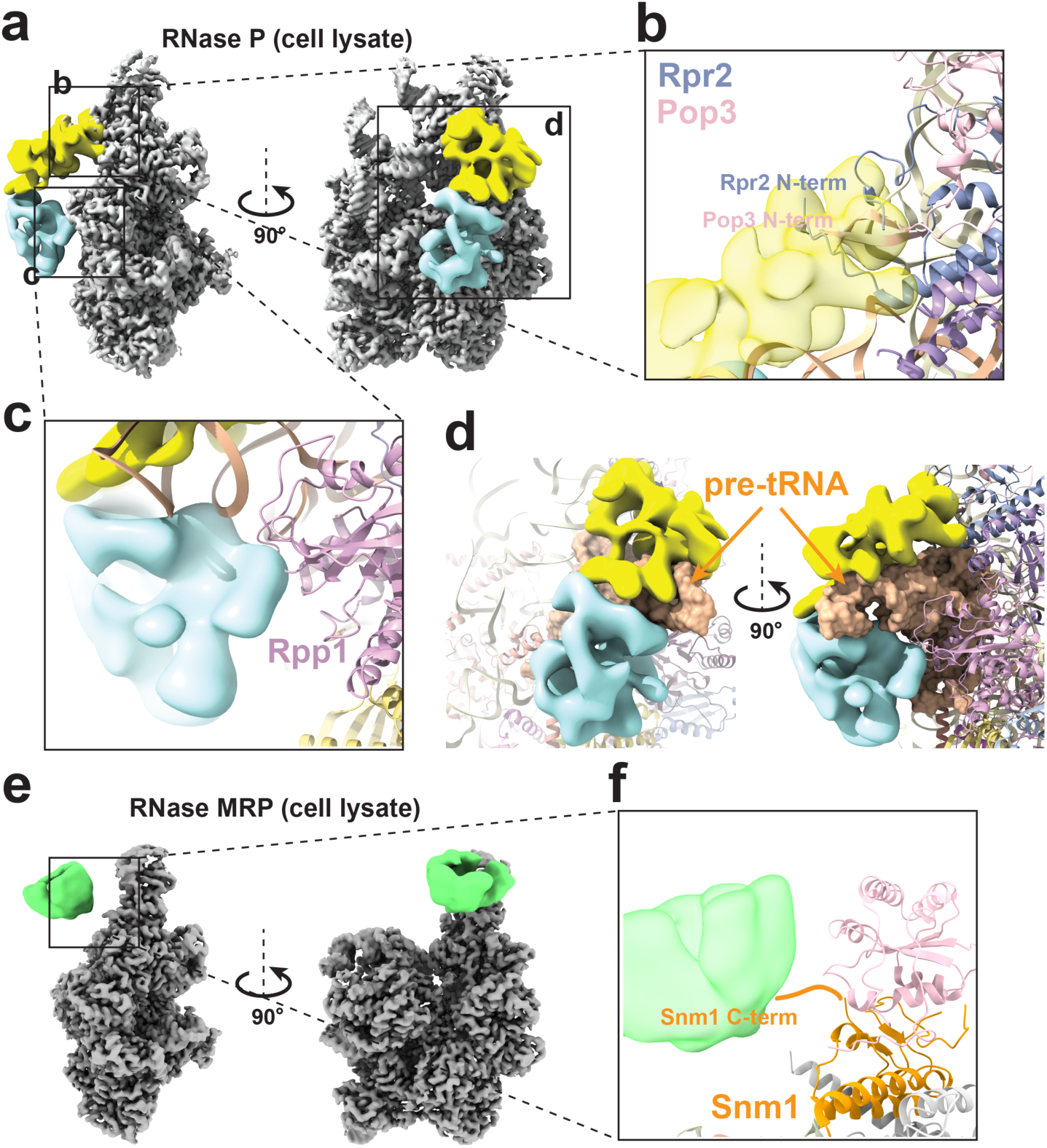
**a,** Cryo-EM structure of RNase P derived from cell lysates, revealing two additional densities (yellow and cyan) identified through multibody refinement. **b,** Close-up view of the upper extra density (yellow), which is connected to the N-terminal regions of Rpr2 and Pop3. **c,** Close-up view of the lower extra density (cyan) located near Rpp1. **d,** Overlay of pre-tRNA (orange; PDB ID: 6AH3) showing that it fits well within the gap between the two extra densities and the core RNase P structure. **e-f,** Cryo-EM structure of RNase MRP from cell lysates after the multibody refinement, highlighting an additional density (green) near the C-terminus of Snm1, a unique component of RNase MRP.

## Discussion

Suppose we apply 5 µl of purified protein solution with a molecular weight of 100 kDa (100,000 g/mol) at a final concentration of 1.0 mg/ml to the cryo-EM grid prior to blotting. If approximately 500 particles are observed per micrograph on a 4k x 6k pixel direct electron detector (1 Å/pixel), calculations indicate that over 99.9% of the protein is lost either to the filter paper or through adsorption onto the amorphous holey carbon (Suppl. Fig. 7). Minimizing this loss could significantly lower the amount of sample needed for cryo-EM analysis.

The graphene-based nanobody-crosslinked affinity grid presented in this study offers significant advancements in cryo-EM specimen preparation by enhancing target specificity, attraction, and population on the grid through affinity tag-probe interactions. Additionally, it mitigates specimen loss during the blotting step by immobilizing the specimens directly onto the grid. This capability addresses critical challenges in the structural determination of endogenous proteins, particularly their low abundance and the presence of contaminants in cell lysates. Moreover, by genetically incorporating the TAP(ATF)-tag into the target gene and expressing the proteins under natural conditions, we can obtain physiologically active and biologically functional proteins with correctly assembled architectures. Here, we demonstrated that the GFD-A grid effectively captures target proteins from both crude cell lysates and anti-FLAG affinity eluates. The grid was shown to not only enrich the target proteins on its surface but also support image acquisition and subsequent data processing, validating its utility for high-resolution cryo-EM structural studies.

The Graffendor grid offers several advantages over other earlier versions of affinity grids. Firstly, Graffendor grids are produced in batch scale (36 grids per batch), ensuring consistent quality within a single batch. Secondly, a genetically modified nanobody is used as the affinity probe, which is covalently attached to the graphene surface via one-step crosslinking. This one-step production process minimizes potential variations in affinity grid production. Additionally, specimens are protected from exposure to the air-water interface by the affinity probe, and the long GS linker (∼100 Å) provides flexibility to potentially alleviate preferred particle orientation in the vitrified ice. Adjusting the length of the GS linker can alter particle orientation thereby changing population of certain 2D projection views. Maintaining the affinity efficacy of the grid during long-term storage provides an additional advantage, ensuring the reliability and reproducibility of experiments over extended periods. This stability allows researchers to prepare and store grids in advance, streamlining workflow and facilitating the structural analysis of target proteins without the need for immediate preparation, thereby improving overall efficiency in cryo-EM studies.

In our proof-of-concept study using *S. cerevisiae* RNase P and RNase MRP with Pop6-ATF as a pull-down component, we unexpectedly observed additional densities in the cryo-EM structures derived from cell lysates at low contour levels. These extra densities, which were absent in structures obtained from purified complexes, may represent transiently associated components of RNase P and RNase MRP (Fig. 6 and Suppl. Fig. 6). Based on their binding locations, these dynamic elements could play a crucial role in substrate recognition and/or enzymatic regulation. This finding highlights an unexpected advantage of the GFD grid in identifying transiently associated components directly from cell lysates.

The use of small volumes of cell culture, combined with a simplified protein purification process, enables the rapid screening of multiple target proteins within a limited timeframe. Notably, AlphaFold-predicted structures have recently become a favored approach for guiding structure-based mutational studies^46^. With a surge in publications presenting AI-generated complex structures lacking empirical validation^47–49^, there is a growing need for tools that facilitate the acquisition of experimentally validated protein complex structures. The Graffendor grid can serve as an effective tool for this purpose. This methodology also facilitates cryo-EM structural studies of proteins that were previously challenging to produce recombinantly or difficult to isolate in sufficient quantities from native sources.

## Method

### Cloning and purification of β-galactosidase-2xALFA, modified NbALFA, modified calmodulin, β-galactosidase-CBP, Cys-mutant of modified NbALFA/modified calmodulin

β-galactosidase-2xALFA, modified NbALFA (with two surface lysine residues mutated to arginines and C-terminally fused to 32 GS-linkers followed by five poly-lysines), modified calmodulin (with five surface lysine residues mutated to arginines and C-terminally fused to 32 GS-linkers followed by five poly-lysines), β-galactosidase-CBP, and Cys-mutants of modified NbALFA (E2C mutation) and modified calmodulin (D3C mutation) were either PCR-amplified or synthesized after codon optimization. These constructs were cloned into the pET3a vector with either His- or His-MBP-tags at the N-terminus. Following transformation into *Escherichia coli* Rosetta (DE3) cells, the proteins were expressed using PA-5052 auto-inducible media^50^. Harvested cells were resuspended in a buffer containing 30 mM Tris-HCl (pH 8.0), 500 mM NaCl, and 3 mM β-mercaptoethanol, supplemented with protease inhibitor cocktails, and sonicated for 2 minutes on ice. The soluble lysate was recovered by centrifugation at 35,000 x g for 1 hour and applied to a cobalt affinity column (Takara) pre-equilibrated with 30 mM Tris-HCl (pH 8.0), 500 mM NaCl, and 3 mM β-mercaptoethanol. The resin was washed with a washing buffer consisting of 30 mM Tris-HCl (pH 8.0), 500 mM NaCl, 3 mM β-mercaptoethanol, and 30 mM imidazole. Target proteins were eluted with an elution buffer containing 30 mM Tris-HCl (pH 8.0), 500 mM NaCl, 3 mM β-mercaptoethanol, and 300 mM imidazole. If the target protein contained a His-MBP tag, the cobalt elution fraction was applied to an amylose resin (New England Biolabs) pre-equilibrated with 30 mM Tris-HCl (pH 8.0), 500 mM NaCl, and 3 mM β-mercaptoethanol. After washing, target proteins were eluted with an amylose elution buffer containing 30 mM Tris-HCl (pH 8.0), 500 mM NaCl, 3 mM β-mercaptoethanol, and 20 mM maltose. Either cobalt elution fractions (His-tag) or amylose elution fractions (His-MBP tag) were dialyzed in 30 mM Tris-HCl (pH 8.0), 500 mM NaCl, and 1 mM DTT at 4 °C overnight with Tobacco Etch Virus (TEV) protease at a 1:100 molar ratio to cut the tag during dialysis. The cleaved tags were removed using additional cobalt and amylose affinity columns. The flow-through fractions were concentrated using an Amicon Centrifugal Filter (Millipore Sigma) and applied to a HiLoad 16/600 Superdex 200 pg size exclusion chromatography column (GE Healthcare) pre-equilibrated with 30 mM HEPES (pH 7.0), 150 mM NaCl, and 1 mM tris(2-carboxyethyl)phosphine (TCEP). Desired fractions were pooled and concentrated for subsequent experiments.

### Making a single batch of the GFD-A grid (total 36 grids)

A total of 36 graphene-coated gold Quantifoil holey carbon grids (Au R1.2/1.3 300 mesh Quantifoil grids, Ted Pella Inc.) were prepared as previously described^30^. The graphene-coated grids were first treated by glow discharge (PELCO easiGlow, Ted Pella Inc.) at 5 mA for 10 seconds under vacuum (< 0.26 mbar). The grids were incubated in 25 mM MES (pH 6.5), 2 mM 1-ethyl-3-(3-dimethylaminopropyl)carbodiimide (EDC) for 5 minutes, followed by the addition of an equal volume of 25 mM MES (pH 6.5), 5 mM Sulfo-N-hydroxysulfosuccinimide (NHS) for 15 minutes at room temperature. The grids were then washed in 25 mM MES (pH 6.5) and incubated in 0.01 mg/mL modified NbALFA (30 mM HEPES [pH 7.0], 150 mM NaCl, 1 mM TCEP) for 2 hours at room temperature with gentle shaking. The resulting GFD-A grids were stored in a glass petri dish containing 30 mM HEPES (pH 7.0), 150 mM NaCl, 1 mM TCEP, and 0.02% (w/v) sodium azide for long-term storage at 4 °C.

### Confocal Microscopy (amylose pulldown/ Cy5 labeling)

To measure the crosslinking efficiency of N-terminal cysteine-incorporated modified calmodulin onto the EDC/Sulfo-NHS–activated surface of the graphene grid, 1 ∼ 2 mg/ml of proteins were degassed in HEPES pH 7.0, 150 mM NaCl, 5 mM TCEP buffer in a plastic vial. Degassing was performed by applying a vacuum to it for several minutes through argon gas. Then, the proteins were labeled with a 20-fold excess amount of sulfo-Cyanine5 maleimide (Cy5, Lumiprobe) overnight at 4°C. Cy5-labeled proteins were immediately purified with a Superdex 200 10/300 SEC column, snap-frozen, and stored at -80°C until use. Fluorescence images of the grids were observed using the Nikon CSU-X1 confocal microscopy (Nikon, Japan) with an excitation peak at 640 nm and an emission peak at 670 nm wavelength.

### Cryo-EM structure determination of β-galactosidase-2xALFA using the GFD-A grid

To evaluate the specificity of the GFD-A grid, either purified β-galactosidase-2xALFA or β-galactosidase-CBP was incubated with the GFD-A grid and imaged using cryo-electron microscopy. A total of 500 µL of 25 μg/ml and 5 μg/ml β-galactosidase-2xALFA or β-galactosidase-CBP was incubated with the GFD-A grid overnight, followed by washing with washing buffer (30 mM HEPES [pH 7.0], 500 mM NaCl, 1 mM TCEP, 0.1% NP-40) overnight at 4 °C with gentle shaking. The GFD-A grids were blotted using a Leica GP2 (5 seconds blotting), and microscopic images were obtained using a 300 KeV Titan Krios equipped with the K3 Summit direct electron detector and the Gatan image filter (± 20 eV) at the University of Michigan cryo-EM facility. Specifically, 1,760 micrographs of the GFD-A grid incubated with 25 μg/ml β-galactosidase-2xALFA were collected for structure determination. A total of 677 K particles were picked using TOPAZ^36^ and a 2.71 Å resolution cryo-EM structure of *β*-galactosidase was determined with 434 K particles using the cryoSPARC software package^37^ (Supp. Table 1).

### In vitro tomography of the GFD-A grid incubated with β-galactosidase-2xALFA

The vitrified GFD-A grid, incubated with 20 μg/mL β-galactosidase-2xALFA and 10 nm gold nanoparticles, was imaged using a 300 kV Titan Krios G4i transmission electron microscope equipped with a K3 Summit direct electron detector and a Gatan image filter. The magnification was set to achieve a pixel size of 2.1 Å (42,000 x magnification) for tilt series data collection. Dose-fractionated images with 3° increment were automatically recorded using SerialEM in counting mode, covering an angular range from –60° to +60° with a dose-symmetric acquisition scheme, resulting in a total dose of 100 e^−^/Å^2^ ^51,52^. The tilt series were processed with IMOD,^53^ where tilt series were aligned using patch tracking and tomograms were reconstructed by weighted back projection, as implemented in IMOD. Tomograms were segmented with the convolutional neural network method implemented in EMAN2.2^54^. Visualization of tomographic volumes was performed using UCSF ChimeraX^55^.

### Budding yeast TAP(ATF)-tagging at the C-terminal Pop6 via homologues recombination

To study structures of Pop6 associated protein complexes, we tagged the C-terminus of this gene with ATF-tag (3x ALFA tag–Tev site–3x flag tag). We used the pFA6 vector to perform PCR with primers specific to the C-terminus of Pop6 for cloning. 120 ml of YPD (Yeast Peptone Dextrose) liquid medium was inoculated with 1.2 ml of *S. cerevisiae* (LWF 7235) at 32 °C and 200rpm until it reached an OD_600_ of 0.5. The 20 ml of cells were centrifuged at 3000 rpm for 3 minutes, and the pellet was resuspended in 1 ml of 1XTE/LiAc (Tris-EDTA/Lithium Acetate) buffer. After resuspension, the cells were centrifuged again under the same condition. The pellet was resuspended in 100 ul of 1X LiAc/TE buffer. A mixture of 15 μl Pop6-ATF PCR product, 6 ul salmon sperm DNA, and 0.7 ml 40% PEG (Polyethylene glycol) was added to the cell suspension. The cells were thoroughly mixed, and heat shocked at 42 °C for 30 minutes. The cells were centrifuged at 3000 rpm for 3 minutes, and the pellet was resuspended in 1ml of YPD medium. The cells were incubated overnight at 32 °C and 200 rpm. The following day, cells were centrifuged again at 3000 rpm for 3 minutes. Finally, the pellet was resuspended in 500 ul of YPD medium. 200 ul of cells were plated on a YPD Hygromycin plate at 30 °C. Yeast transformants were counted after 2 days of growth.

### Pop6-ATF yeast strain cell culture and preparation of cell lysate and anti-flag eluate for the GFD-A grid incubation

One liter of Pop6-ATF yeast cells were harvested at OD_600_ 2.0 and frozen as ice balls with liquid nitrogen. The cells were broken using a 6875D Freezer Mill (SPEX Sample Prep.). The cells were resuspended in 50 ml of cell lysis buffer (30 mM HEPES [pH 7.0], 500 mM NaCl, 5% glycerol, 0.1% NP40, 1 mM TCEP) with protease inhibitor cocktail and sonicated. The cells were centrifuged at 18,000 rpm for 1 hour at 4 °C, repeated twice. The supernatant was collected and filtered with 5.0 um syringe filters (Sartorius Minisart NML). The 50 ml of supernatant was transferred into a column containing anti-FLAG M2 beads (Sigma-Aldrich). Subsequently, the beads were washed with 10 ml of washing buffer (30 mM HEPES [pH 7.0], 500 mM NaCl, 5% glycerol, 0.1% NP40). Finally, the beads were incubated overnight with TEV protease in the elute buffer (30 mM HEPES [pH 7.0], 150 mM NaCl, 5% glycerol, 0.1% NP40). The flag beads were centrifuged at 800 rpm for 5 minutes at 4 °C to collect the protein in the supernatant.

### Cryo-EM structure determination of RNase P and RNase MRP using cell lysate/GFD-A and anti-flag eluate/GFD-A

Both cell lysate/GFD-A and anti-flag eluate/GFD-A grids were plunge-frozen using a Leica GP2 (5 second blotting; 95% humidity), and their micrographs were collected using a 300 keV Titan Krios transmission electron microscope equipped with a K3 Summit direct electron detector and a Gatan image filter (± 20 eV) at the University of Michigan cryo-EM facility. A total of 4,924 micrographs and 5,148 micrographs were collected, and 1.17 million particles and 1.1 million particles were picked and extracted for cell lysate/GFD-A and anti-flag eluate/GFD-A grids using TOPAZ^36^, respectively. After extensive 2D and 3D classification and 3D refinement using the cryoSPARC^37^ and RELION software suites^56^, final cryo-EM maps were obtained with resolutions of 3.0 Å (RNase P) and 3.3 Å (RNase MRP) from cell lysate/GFD-A, and 3.9 Å (RNase P) and 3.6 Å (RNase MRP) from anti-flag eluate based on the FSC=0.143 gold standard, respectively.

## Acknowledgments

We thank Drs. Vinson Lam, Ashleigh Raczkowski, and Alexandrea Rizo at the University of Michigan cryo-EM facility for assistance with data collection. This work was supported by grants (CA 250329, NS 116008, and AI 160067) to U.S.C. A patent application related to aspects of this research has been filed (U.S. patent Application No. 18/833,720).

## Author Contributions

S.A. purified proteins, performed assays, and revised the manuscript, E.A. conceptualized and revised the manuscript. T.K. purified proteins and performed experiments. S.P. purified proteins and performed experiments. B.S. purified proteins and performed experiments. K.P.R. generated yeast strains, purified proteins, and performed experiments. G.L. purified proteins and performed experiments. C.K. conceptualized and revised the manuscript. B.K. performed experiments. H.K. performed experiments. S.P. performed experiments. D.T. assisted and consulted in the cryo-EM structure determination. U.S.C. conceptualized, generated the GFD-A grids, prepared cryo-EM samples, determined the cryo-EM structure, analyzed data, and wrote the manuscript.

## Data availability

The accession numbers for β-galactosidase-2xALFA, *S. cerevisiae* RNase MRP (anti-flag eulate), RNase P (anti-flag eulate), RNase MRP (cell lysate), and RNase P (cell lysate) cryo-EM maps are EMD-48651, 48652, 48653, 48652, 48653, respectively.

## Competing interests

The authors declare no competing interests.

*Corresponding Author; Tel.: 1-(734) 764-6765; Fax: 1-(734) 763-4581.

