## Supplementary data for "The graphene-based affinity cryo-EM grid for the endogenous protein structure determination"

**Supplementary Table 1. Cryo-EM Data Collection, Refinement, and Validation Statistics**

| | β-galactosidase-2xALFA | RNase P (cell lysate) | RNase MRP (cell lysate) | RNase P (anti-flag eluate) | RNase MRP (anti-flag eluate) |
|---|---|---|---|---|---|
| **Data Collection and Processing** | | | | | |
| Magnification | 81,000 | 81,000 | | 81,000 | |
| Voltage (kV) | 300 | 300 | | 300 | |
| Electron exposure (e-/Å$^2$) | 66 | 60.9 | | 60 | |
| Defocus range (μm) | -0.5 to -2.0 | -1.0 to -2.5 | | -0.8 to -2.0 | |
| Pixel size (Å) | 1.08 | 1.08 | | 1.08 | |
| Symmetry imposed | D2 | C1 | | C1 | |
| Number of micrographs collected | 1,761 | 4,924 | | 4,681 | |
| Initial particle images (no.) | 676,763 | 1,170,230 | | 1,107,285 | |
| Final particle images (no.) | 438,825 | 293,742 | 268,844 | 44,258 | 130,068 |
| Map resolution (Å) | 2.71 | 3.0 | 3.3 | 3.9 | 3.6 |
| FSC threshold | 0.143 | 0.143 | 0.143 | 0.143 | 0.143 |
| EMD | EMD-48651 | EMD-49177 | EMD-49178 | EMD-48653 | EMD-48652 |

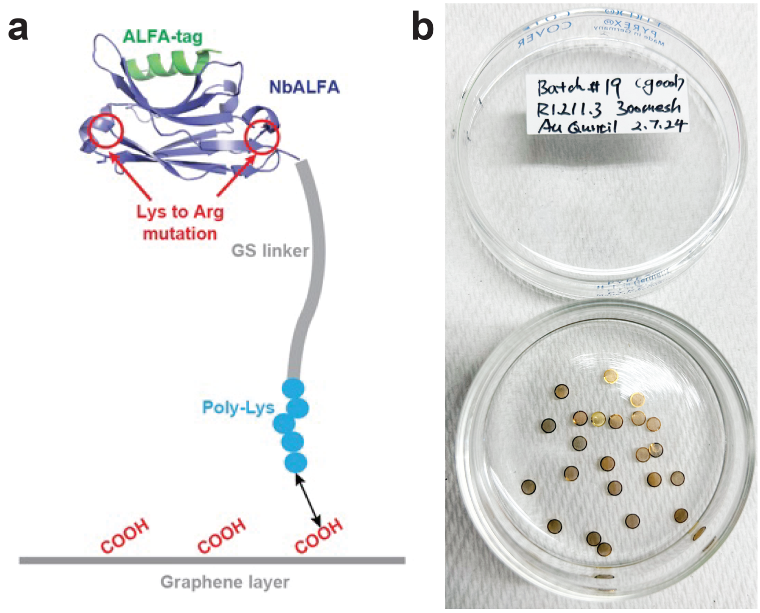


**Supplementary Figure 1. a,** Schematic representation of the modified NbALFA as an affinity probe. Five lysine residues at the C-terminal end of modified NbALFA (cyan circles) facilitate crosslinking with carboxyl groups on the oxidized graphene surface. **b,** Production and storage of a single batch of GFD-A grids. A total of 36 GFD-A grids are generated from a single batch of graphene grids, with all undergoing modified NbALFA crosslinking simultaneously to ensure consistent quality. Following production, the GFD-A grids are stored in a glass petri dish containing storage buffer (30 mM HEPES, pH 7.0, 150 mM NaCl, 0.1% NP-40, and 0.02% $NaN_3$).

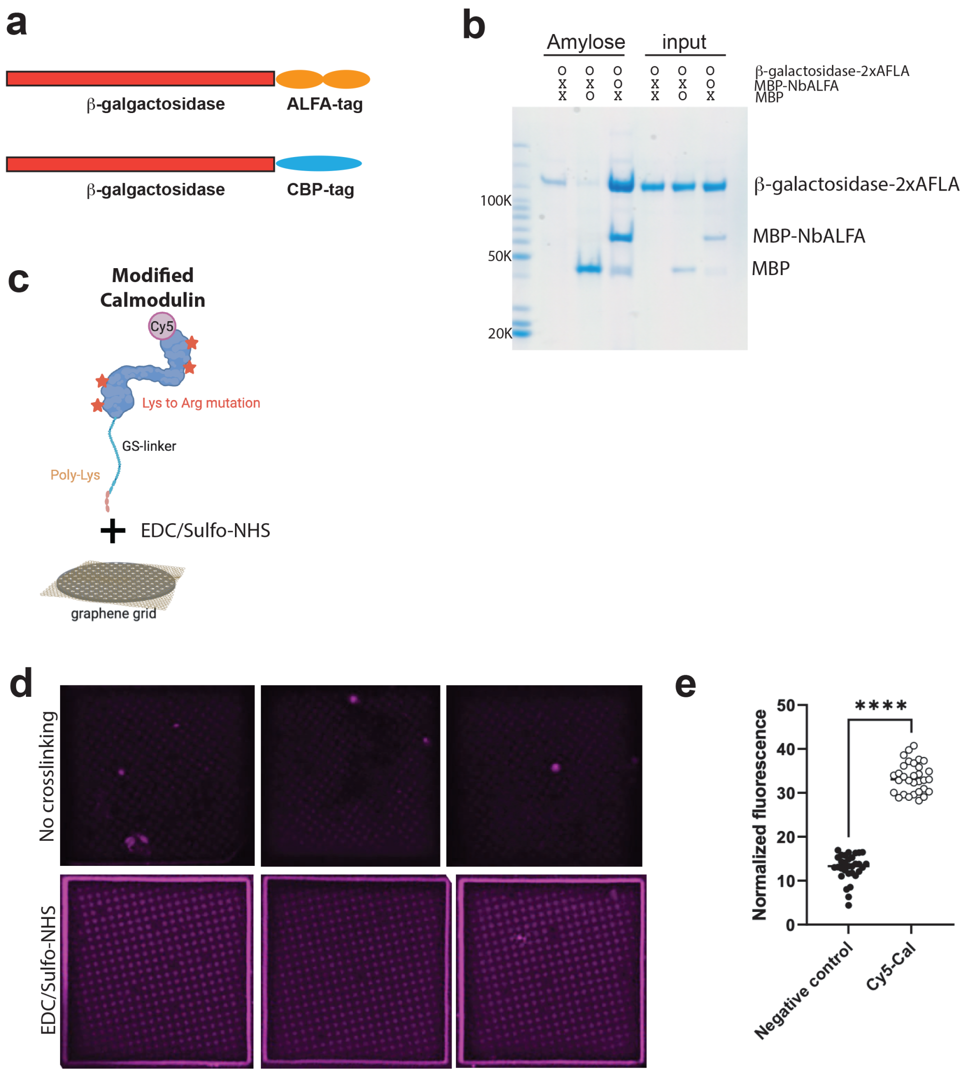


**Supplementary Figure 2. a,** Schematic representation of the β-galactosidase-2xALFA and β-galactosidase-CBP constructs used in this study. **b,** Amylose pull-down assay of MBP-modified NbALFA with β-galactosidase-ALFA.tag. **c,** Diagram illustrating the crosslinking of Cy5-labeled modified calmodulin onto the oxidized graphene grid surface using EDC/Sulfo-NHS. **d-e,** Confocal microscopy images of Cy5-labeled modified calmodulin on the graphene grid surface, either without a crosslinking reagent (top) or with EDC/Sulfo-NHS treatment (bottom). We measured the fluorescence intensity of each hole from 10 holes on 3 different grids for both the negative control and Cy5-labeled samples, and normalized the values using the Fiji program. $P$ value was calculated using the Student's $t$-test (**** means $P$<0.0001).

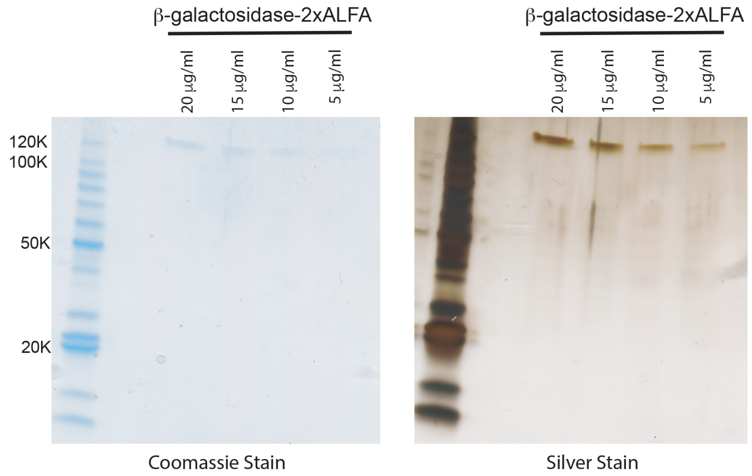


**Supplementary Figure 3.** The Coomassie and silver staining of the SDS-PAGE gel run with different concentrations (20, 15, 10, 5 μg/ml) of β-galactosidase-2xALFA.

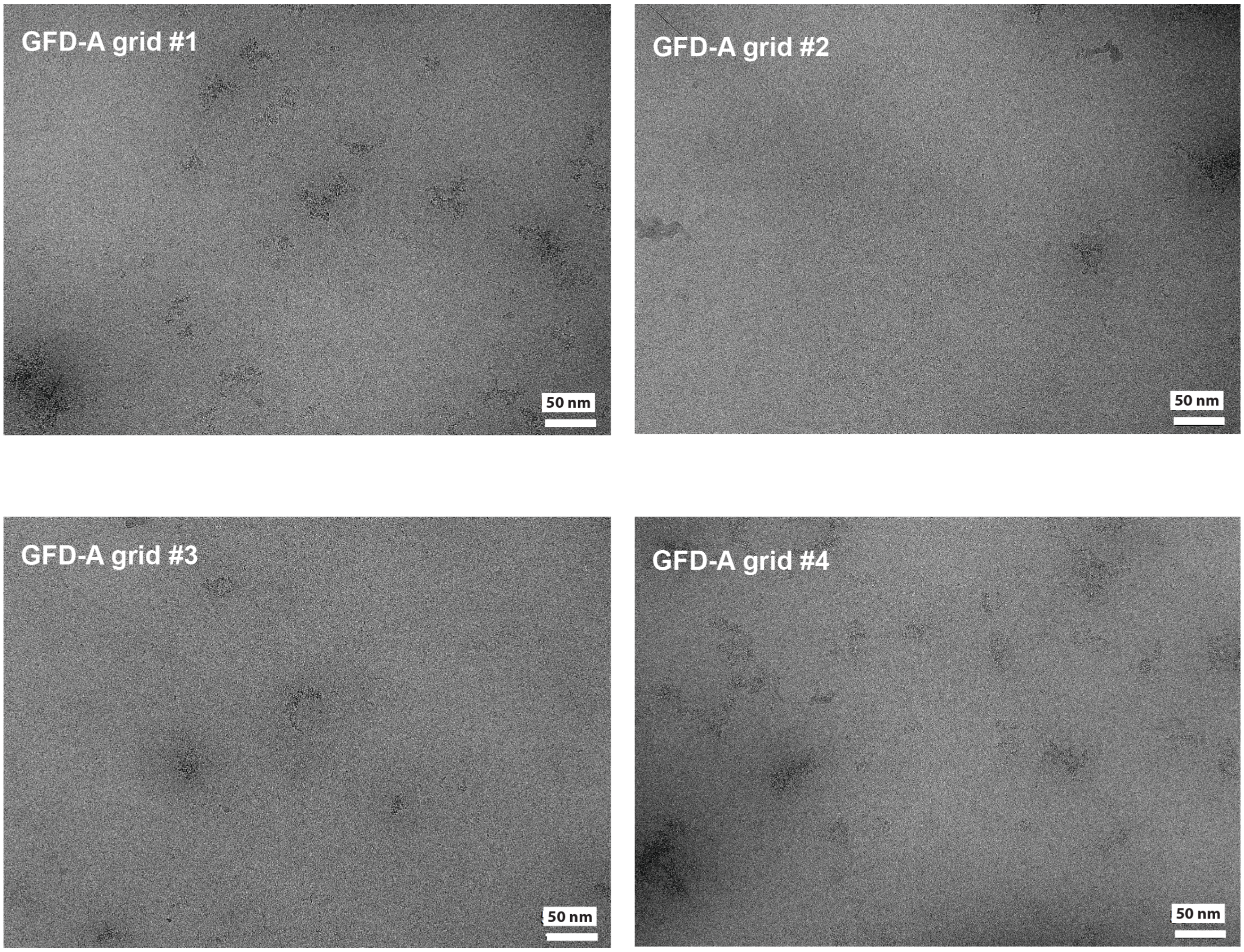


**Supplementary Figure 4.** Micrographic images of four GFD-A grid replicates within the same batch. Although a certain level of PMMA contaminants is observed (Ahn 2023){Ahn, 2023 #78}, it does not interfere with the image acquisition, particle picking, and data processing.

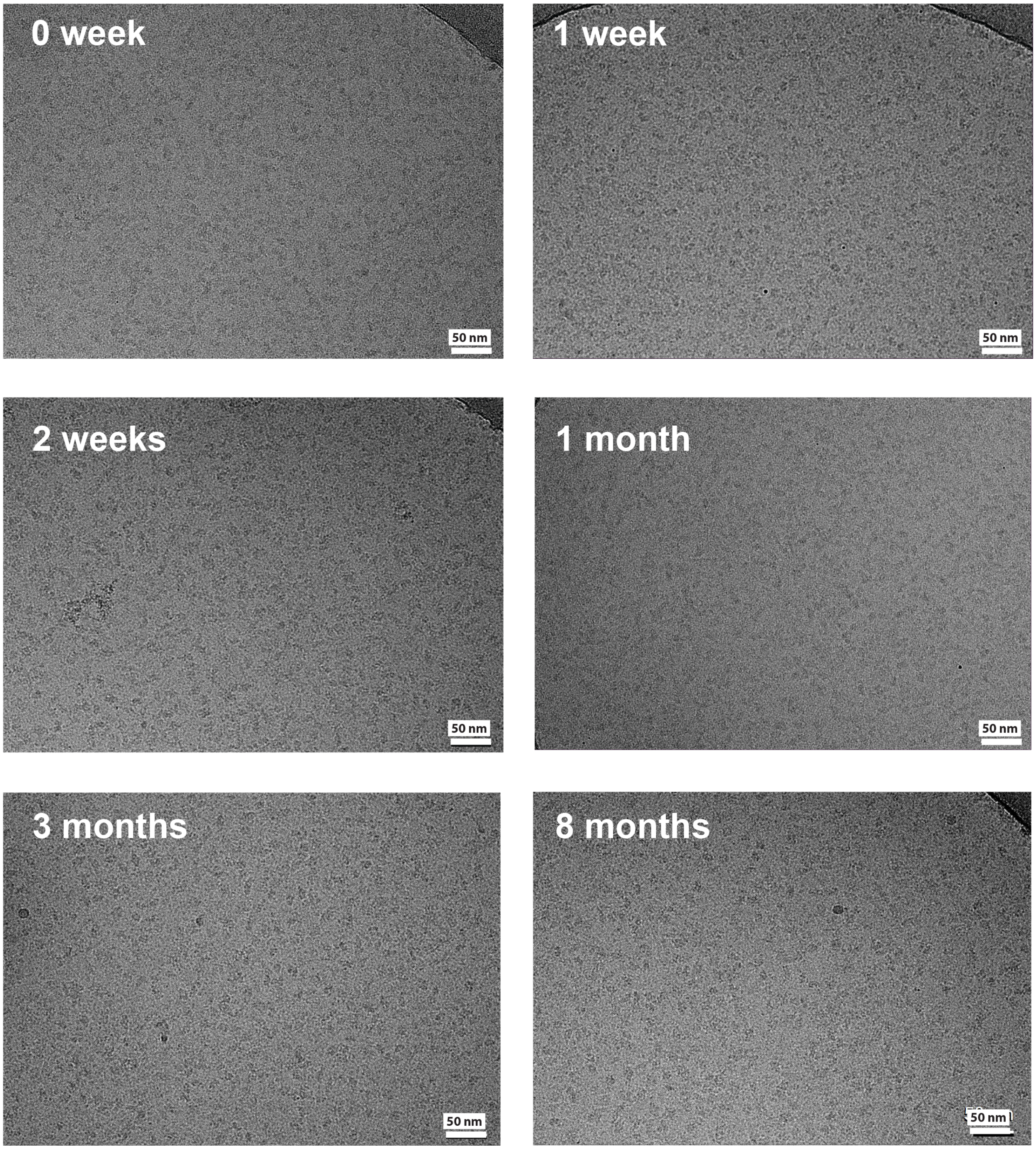


**Supplementary Figure 5. Long-term storage of the GFD-A grid in the storage buffer.** The potential quality change of the GFD-A grid stored in the storage buffer (30 mM HEPES, pH 7.0, 150 mM NaCl, 0.1% NP-40, and 0.02% $NaN_3$; 4°C) was assessed by acquiring micrographs of β-galactosidase-2xALFA/GFD-A grids at various time points (0, 1, and 2 weeks; 1, 3, and 8 months). The GFD-A grid remained stable for at least 8 months under these storage conditions, indicating its suitability for long-term preservation.

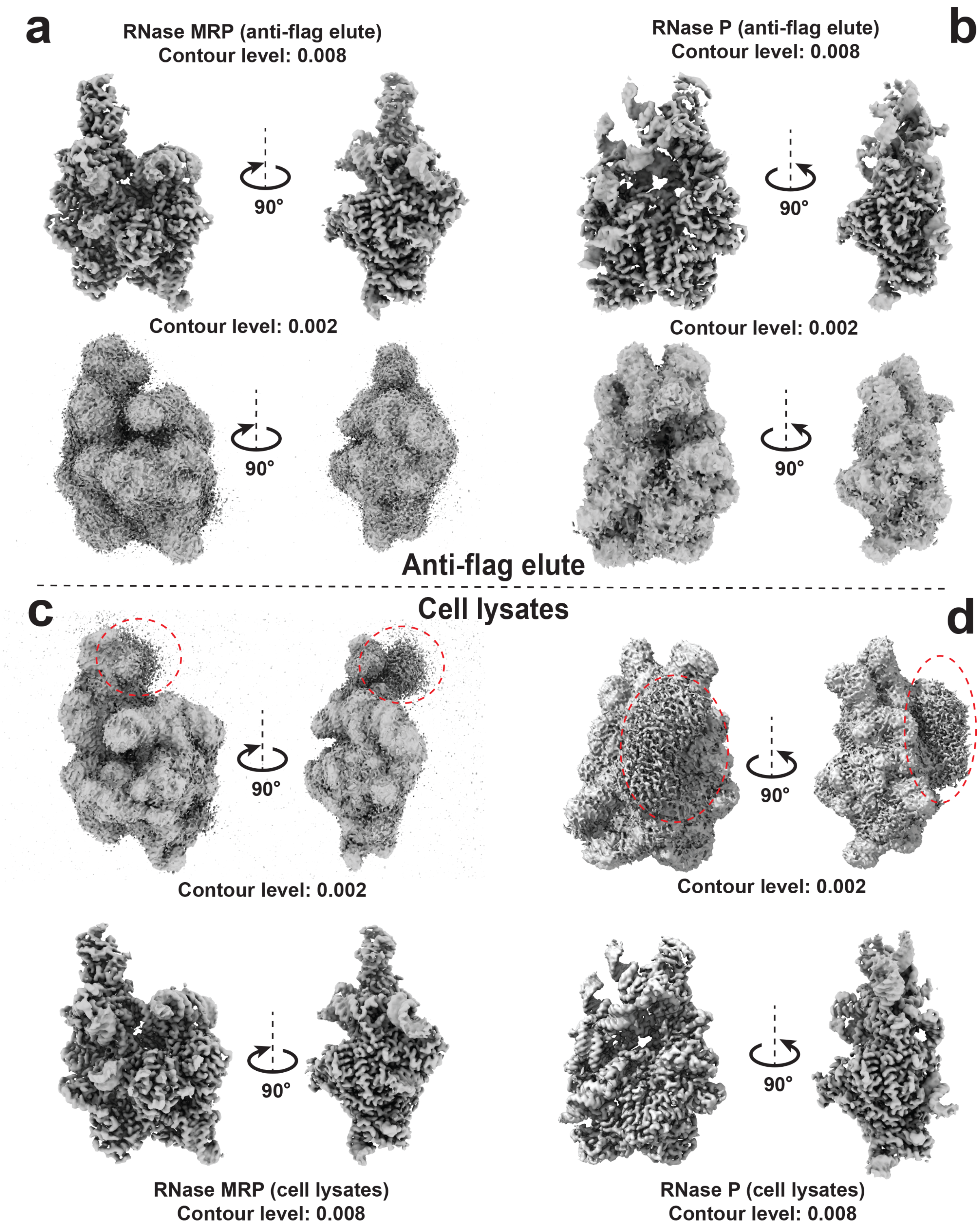


**Supplementary Figure 6. Identification of extra densities in cryo-EM maps of RNase P/MRP derived from cell lysates at low contour levels. a-d,** Cryo-EM structures of RNase P/MRP obtained from anti-FLAG eluate and cell lysate. Maps are shown at two different contour levels: 0.008 and 0.002. Notably, extra densities were exclusively observed in the cryo-EM maps of RNase P/MRP derived from cell lysates at the contour level of 0.002 (indicated by red dashed circles).

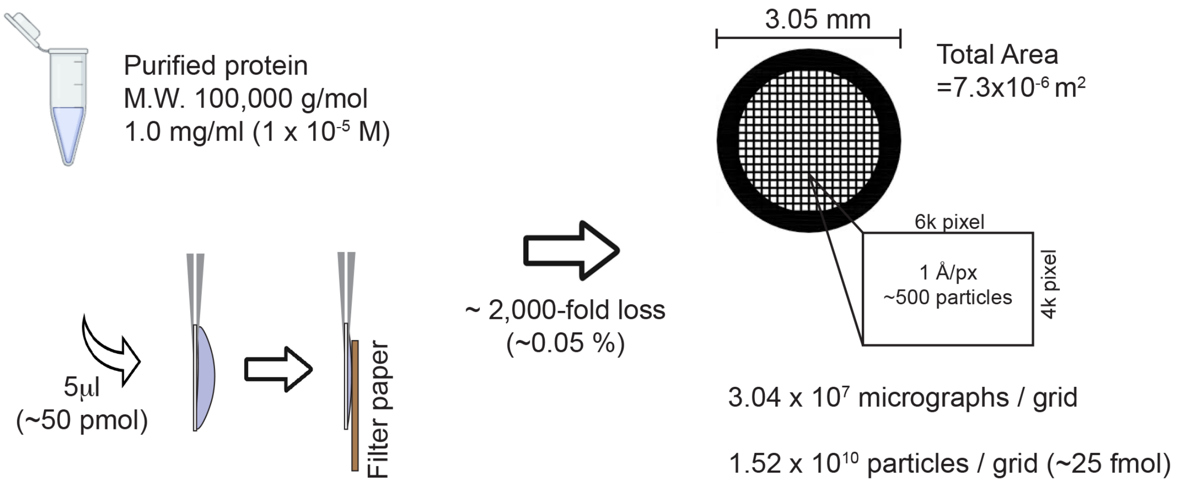


**Supplementary Figure 7. Estimation of protein loss during cryo-EM grid preparation.** For a purified target protein with a molecular weight of 100 kDa at a concentration of 1.0 mg/ml, applying 5 µl onto the cryo-EM grid corresponds to approximately 50 pmol of protein. Based on an average of ~500 particles per 4k × 6k micrograph at 1.0 Å/pixel resolution, the estimated total number of particles retained on the grid is approximately 25 fmol. This analysis underscores the substantial protein loss during blotting, highlighting the necessity for improved methods to enhance particle retention and minimize sample wastage.
